# Evolutionary rescue in one dimensional stepping stone models

**DOI:** 10.1101/2020.10.29.360842

**Authors:** Matteo Tomasini, Stephan Peischl

## Abstract

Genetic variation and population sizes are critical factors for successful adaptation to novel environmental conditions. Gene flow between sub-populations is a potent mechanism to provide such variation and can hence facilitate adaptation, for instance by increasing genetic variation or via the introduction of beneficial variants. On the other hand, if gene flow between different habitats is too strong, locally beneficial alleles may not be able to establish permanently. In the context of evolutionary rescue, intermediate levels of gene flow are therefore often optimal for maximizing a species chance for survival in metapopulations without spatial structure. To which extent and under which conditions gene flow facilitates or hinders evolutionary rescue in spatially structured populations remains unresolved. We address this question by studying the differences between evolutionary rescue in the island model and in the stepping stone model in a gradually deteriorating habitat. We show that evolutionary rescue is modulated by the rate of gene flow between different habitats, which in turn depends strongly on the spatial structure and the pattern of environmental deterioration. We use these insights to show that in many cases spatially structured models can be translated into a simpler island model using an appropriately scaled effective migration rate.

## Introduction

Recent environmental or climatic changes have severe consequences on the survival of many species [Thomas et al., 2004]. Rapid and severe environmental change might cause extinction of resident populations due to genetic maladaptation to the new environmental conditions [Cahill et al., 2013]. A population may survive such environmental change if it adapts quickly enough via standing genetic variation or *de novo* mutations, a process that has been coined evolutionary rescue [Gomulkiewicz and Holt, 1995, Alexander et al., 2014, Carlson et al., 2014]. Theoretical studies of the mechanisms that can facilitate or hinder survival of populations experiencing environmental change can provide critical insights in the context of adaption and survival in the face of global climatic change, but also in medically relevant scenarios such as the evolution of antibiotic or other treatment resistance (*e.g*. Alexander et al. [2014], Carlson et al. [2014]).

It has been demonstrated theoretically as well as experimentally that evolutionary rescue in single, isolated populations depends strongly on two factors: initial population size and standing genetic variation [Lynch, 1993, Bell and Gonzalez, 2009, Samani and Bell, 2010, Bell and Gonzalez, 2011, Bell, 2013, Ramsayer et al., 2013, Carlson et al., 2014]. In large populations or in populations with high mutation rates, mutations that ultimately rescue a population have higher chances to arise and establish before the population goes extinct [Lynch, 1993, Gomulkiewicz and Holt, 1995, Bell, 2013]. Similarly, initially high standing genetic variation increases the chance that a mutation that can save the population may already be present in the population [Gomulkiewicz and Holt, 1995, Barrett and Schluter, 2008, Vander Wal et al., 2013]. Other factors, such as the competition between individuals, the rate of environmental decay or gene flow from other populations can have more subtle effects and affect the chance of rescue positively or negatively [Uecker et al., 2014, Marrec and Bitbol, 2020].

Evolutionary rescue has been thoroughly studied in idealized laboratory conditions and key theoretical predictions have been confirmed [Carlson et al., 2014]. However, these idealized laboratory conditions are unlikely to reflect the spatio-temporal demographics of natural populations or the intricate spatial and population structure within and across human bodies during the evolution of antibiotic or treatment resistance [Bell, 2017]. For instance, in the face of structured populations it has been demonstrated both experimentally [Bell and Gonzalez, 2009, 2011, Gonzalez and Bell, 2013] and theoretically [Uecker et al.,2014, Czuppon et al., 2021] that gene flow between sub-populations can facilitate evolutionary rescue. The benefits of spatial structures and dispersal to adaptation and evolutionary rescue have also been demonstrated in other contexts. For example, dispersal can help to spread beneficial mutations causing rapid population differentiation [Feder et al., 2019], and heterogeneity in environmental conditions caused by imperfect drug penetration allowed the evolution of multi-drug resistance [Moreno-Gamez et al., 2015]. In the context of antibiotic resistance evolution, interesting examples come from *Pseudomonas aeruginosa* where the formation of biofilms are hallmarks of disease progression and chronic infections [Botelho et al.,2019]. Experimental studies have shown that the spatial structure and environmental heterogeneity in biofilms may also contribute to an increased likelihood of resistance evolution as compared to well-mixed populations [Ahmed et al., 2018].

Uecker et al. [2014] significantly improved our understanding of how dispersal can help beneficial mutations to cause evolutionary rescue in an island model, while subsequent work about the benefits of dispersal was done by Tomasini and Peischl [2020] for a two-deme model. A critical assumption of these modelling attempts is that gene flow occurs at the same rate between any pair of demes. Experimental and theoretical work suggests, however, that the spatial organization of sub-populations, as well as biased dispersal, modulate the effect of gene flow [Bell and Gonzalez, 2011, Gonzalez and Bell, 2013, Czuppon et al., 2021]. Whether and how spatial structure might change our understanding of the role of gene flow during adaptation to severe environmental change remains an unresolved question.

In this work we extend the work of Uecker et al. [2014] (in an island model) and Tomasini and Peischl [2020] to study the role of gene flow on evolutionary rescue in spatially extended populations using stochastic simulations. In particular, we contrast two extreme cases of gene flow: long-range migration (*i.e*. the island model [Wright, 1943], where migrants are exchanged among all demes at equal rates) and short-range migration (*i.e*. the stepping stone model [Kimura and Weiss, 1964], where migrants move only between neighboring demes).

We find that short- and long-range dispersal can both have a beneficial effect on evolutionary rescue, and that gene flow is beneficial for evolutionary rescue under quite similar conditions in the two models. More precisely, under certain conditions, we find that the island model and the stepping stone model appear equivalent if we consider an effective migration rate that quantifies migration between deteriorated and non-deteriorated areas of the environment. If rescue mutations are strongly deleterious in one environment but not (close to) lethal, we find however qualitative differences between the two models. Our findings suggest that the number of individuals flowing between deteriorated and non-deteriorated parts of the environment is a key quantity in spatially or otherwise structured populations, rather than the rate of gene flow between demes.

## Materials and Methods

### Model

For this study, we take inspiration and expand the work done by Uecker et al. [2014] for the island model to the stepping stone model. We consider a structured population consisting of *D* demes organized along a one-dimensional lattice (see figure 1). Demes are labeled *i* = 1,…, *D* and are filled with haploid wild-type individuals with discrete and non-overlapping generations. Each deme can be in one of two environmental states: the original unperturbed state (or “old” state) and the deteriorated state (or “new” state). In our model, initially the whole habitat is at demographic equilibrium in the original environment. We assume deterioration from left to right (see figure 1) although the model is independent of the direction of deterioration. Every *θ* generations the leftmost deme in the unperturbed state switches to the deteriorated state such that the whole habitat gradually deteriorates at a constant rate of one deme each *θ* generations. In the deteriorated part of the environment, growth rates of wild-type individuals are negative but beneficial mutations can arise and save the population from extinction by restoring positive growth rates.

**Figure 1:**
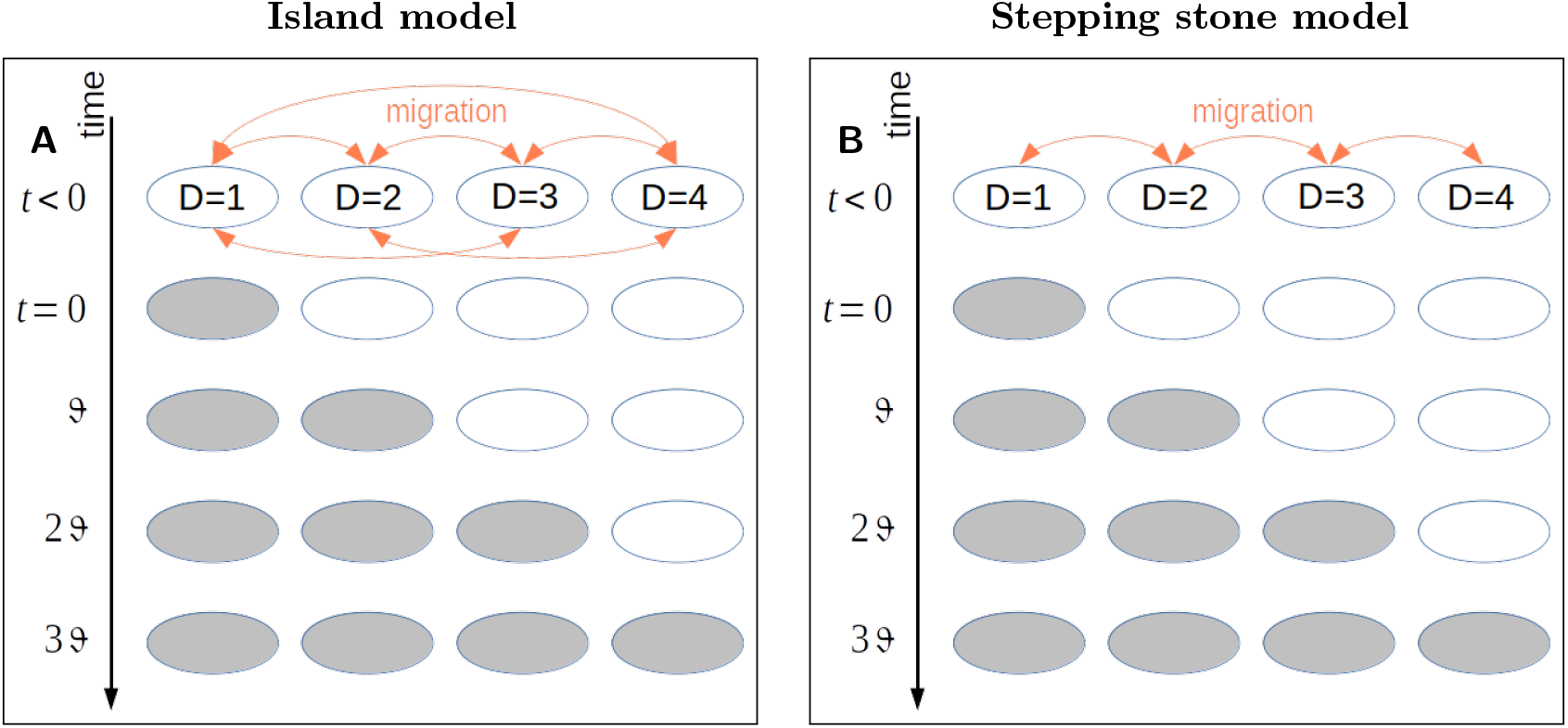
In figure A, a schematic of the island model (global migration), in figure B a schematic of the stepping stone model (local migration). In time, demes become deteriorated (gray demes) one after the other each *θ* generations, while the others remain untouched (white demes). Schematic for *D* = 4.

Population size in deme *i* at time *t* is denoted *N_i_*(*t*) (*i* ∈ {1,…, *D*}, *t* = 0, 1, 2,…). In the original environment, density regulation is implemented such that all demes are at carrying capacity, *κ*, at the end of each generation. This model of instantaneous growth or culling is a good approximation to logistic density regulation if growth rates are sufficiently large relative to migration rates [Beverton and Holt, 1957, Tomasini and Peischl, 2020]. Initially all demes are at carrying capacity such that *N_i_*(0) = *κ*, for all *i*. We define the total carrying capacity of the habitat as *K*_tot_ = *Dκ*. We set *K*_tot_ constant through the rest of the paper, as specified in table 1 (but see [Carlson et al., 2014] for more comprehensive studies of evolutionary rescue for different population sizes). This means that varying *D* corresponds to varying *κ* (as the two are inversely proportional). We call *m* the forward migration rate, that is the proportion of individuals in each deme that migrate [Karlin, 1982, Vuilleumier et al., 2010]. We consider two modes of migration: (i) global migration, allowing an individual to migrate to any deme in the habitat (the island model, Wright [1943]; figure 1A), and (ii) local migration, that allows an individual to migrate only between neighboring demes (the stepping stone model, Kimura and Weiss [1964]; figure 1B). The boundaries of the habitat (to the left of the first deme and to the right of the last deme) are reflecting, that is, individuals cannot leave the habitat.

**Table 1:** List and description of all parameters.

| Parameter | Description |
| --- | --- |
| $D$ | Number of demes |
| $N_i(t)$ | Number of individuals in deme $i$ |
| $\kappa$ | Carrying capacity per deme |
| $K_{\text{tot}} = \kappa \cdot D$ | Total carrying capacity of the habitat, fixed at $2 \times 10^4$ through the paper |
| $m$ | Rate of migration per individual |
| $w_{\text{wt}}^{(o)} = 1$ | Relative fitness of a wild-type individual in the old environment |
| $w_{\text{wt}}^{(n)} = 1 - r$ | Relative fitness of a wild-type individual in the new environment |
| $w_{\text{mu}}^{(o)} = 1 - s$ | Relative fitness of a mutant individual in the old environment |
| $w_{\text{mu}}^{(n)} = 1 + z$ | Relative fitness of a mutant individual in the new environment |
| $0 < s \leq 1$ | Disadvantage against a mutant copy in the old environment |
| $0 < z \ll 1$ | Selective coefficient of a mutant copy in the new environment |
| $0 < r < 1$ | Stress against the wild-type population in the new environment |
| $\theta$ | Speed of deterioration of the environment, length of an epoch |
| $\Theta = \theta(D - 1)$ | Deterioration time of the whole habitat |
| $u = 1/K_{\text{tot}}$ | Mutation probability (wild type to mutant) |

Relative fitness is determined by a single locus with two alleles: a wild-type allele and a mutant allele. We assume that each individual has Poisson-distributed offspring with the mean proportional to its fitness. In the original environment, fitness of wild-type individuals is set to 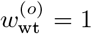 and the fitness of a mutant is 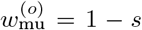, where 0 < *s* < 1 is the selection coefficient against the mutant. Mutations are therefore counter-selected in the original habitat. Absolute fitness in the original habitat is then given by 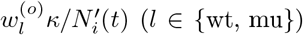, where 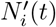 is the population size after selection and migration but before density regulation. Mutations occur with probability *u* per generation and we ignore back mutations. Previous work based a beneficial mutation at a single biallelic locus has shown that the probability of rescue depends monotonically on the product of population size and mutation rate [Orr and Unckless, 2008]; for the rest of this work, we set *u* = 1/*K*_tot_ such that on average a single mutation occurs per generation in the whole habitat, without loss of generality. Because our focus is to study the interaction of gene flow, spatial structure and demographic dynamics on evolutionary rescue, we choose a simple genetic architecture that allows us to investigate our model in detail without having to account for clonal interference [Gerrish and Lenski, 1998], the distribution of fitness effects [Anciaux et al., 2018], or recombination and multi-step rescue [Uecker and Hermisson, 2016, Osmond et al., 2020].

At a time *t* = 0, the first (*i.e*. leftmost in figure 1) deme (*i* = 1) switches its environmental state to the deteriorated environment. In a deteriorated deme, wild-type individuals have an absolute fitness of 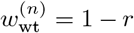, with 0 < *r* < 1, and the resident population starts to decline. In the new environment, the mutant has an absolute fitness 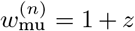, with *z* > 0. We generally chose a value of *z* large enough to just restore positive growth after deterioration. The relative fitness of mutants is then (1 + *z*)/(1 − *r*) = 1 + (*r* + *z*)/(1 − *r*). In the range of parameters that we chose, (*r* + *z*)/(1 − *r*) is often larger than the cost of having the mutation in the non-deteriorated area, *s*. Rescue mutations generally have small deleterious effects before environmental change [Melnyk et al., 2015]. However, recent examples from natural populations show that selection coefficients of rescue mutations can be very large, including mutations that lead to a 5-fold survival increase in females but are lethal in males [Campbell-Staton et al., 2021]. At time *t* = *θ*, deme *i* = 2 (at that point the leftmost deme in the old state) deteriorates, at *t* = 2*θ* the third deme, and so on until the whole habitat is deteriorated. We call the period between two subsequent deterioration events an epoch and *θ* is the length of an epoch. The deterioration of the whole habitat takes Θ = (*D* − 1)*θ* generations (see table 1 for a full list of all parameters of the model).

### Simulations

We performed stochastic simulations. At the beginning of the simulation, *D* demes are filled each with *κ* individuals (*K*_tot_ = *κD* = 2 × 10^4^). Before any deterioration occurs, we leave a burn-in time of 500 generations to generate standing genetic variation.

#### Reproduction

We implement reproduction and selection by sampling from a Poisson distribution with the mean of the distribution given by the relative fitness of wild-type and mutant individuals. Thus the number of wild-type and mutant individuals (here simplified with the superscript *l* ∈ {wt, mu}) changes as

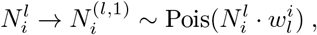

where the superscript ^(1)^ represents the first step of the generation (reproduction). For simplicity, in this section we used 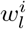 as opposed to 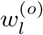 or 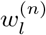, as the fitness depends on the state of deme *i* – *e.g.* at time 0 < *t* < *θ*, 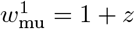 and 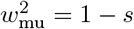.

#### Mutation

After reproduction, we implement mutation as a binomial process with sampling probability *π* = *u*:

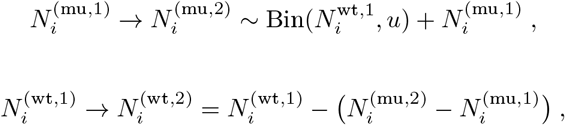

where the superscript ^(2)^ represents the second step of the generation (mutation). For wildtypes, the term 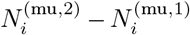 is simply the number of new mutants generated via binomial sampling (and thus removed from the wildtype pool).

#### Migration in the island model

In the island model, the total number of migrants is determined in each deme with a binomial process with sampling probability *π* = *m*, then the whole pool of migrants is re-distributed over all demes with a multinomial distribution. The process is done in the same way separately for mutants and wild types, but for simplicity we will skip the type subscript *l*. For each deme *i*, the number of migrants 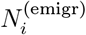 is given by

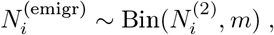

and the total number of individuals in the pool is

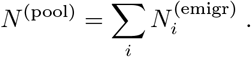

The individuals in the migrant pool, *N*^(pool)^, are then redistributed multinomially, with each individual having the probability 1/*D* of migrating to each deme.

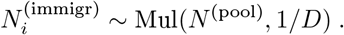

Finally, for each type *l* (superscript omitted), the number of individuals in each deme after migration is

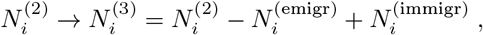

where the superscript ^(3)^ represents the third step in the generation (migration).

#### Migration in the stepping stone model

In the stepping stone model, individuals migrating to the right and individuals migrating to the left are picked as a binomial process with probability *π* = *m*/2 (superscript *l* omitted):

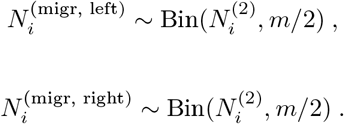

Finally, the number of individuals in deme *i* after migration for each type is updated as

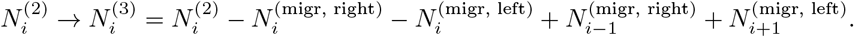

Note that in our formulation the walls of the habitat are reflecting, thus in the first deme 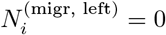 and in the last deme 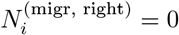.

#### Density regulation

The last phase of the life cylce is density regulation, and it only occurs in non-deteriorated demes. As density is kept constant in the original environment, we define a new population of *K_i_* individuals, where *K_i_* ~ Pois(*κ*), and we sample the number of mutants and wildtypes according to the frequencies after migration, *N*^(1,3)^/(*N*^(mu,3)^ + *N*^(wt,3)^).

#### Burn-out phase

After the whole habitat is deteriorated, we let the system evolve for 500 more generations. The expected extinction time in the absence of evolutionary rescue is much shorter than this burn-off. We consider the population rescued if the mutant population reaches *K*_tot_/2.

We performed 1000 replicates for each parameter combination. Simulation were written in Python and the code can be found on the authors Github page. To test the accuracy of our simulations, we compared them against known cases (see Appendix B). Our simulation recovers expected results for the case with *m* = 0 (figure S2), *θ* = 0 (figure S3) and *D* = 2 (figure S4).

## Results

### Optimal migration rate depends on cost of rescue mutations

From previous work on island models we know that there are three different effects of migration on evolutionary rescue. First, migration provides a demographic rescue effect where gene flow from the original habitat delays extinction in the deteriorated habitat, thus increasing the probability for rescue mutations to occur and spread in the deteriorated habitat. For larger migration rates, rescue mutations are unlikely to establish because they are removed from demes where they are beneficial [Tomasini and Peischl, 2018] (*i.e*., gene swamping occurs [Bulmer, 1972, Lenormand, 2002]). Finally, when migration rates are very large, migration will deplete the original habitat such that density regulation leads to higher absolute fitness in the original habitats, which increases the establishment probability of rescue mutations [Tomasini and Peischl, 2018]. This effect has been called relaxed competition [Uecker et al.,2014] and is especially potent if the fitness cost of rescue mutations is low and if growth rates are high. Figure 2 shows the probability of rescue as a function of the migration rate for different combinations of selection intensities and varying numbers of demes. We identify three qualitatively different scenarios, depending on the strength of selection against rescue mutations in unperturbed environments. If the cost of the mutation in the original environment is relatively low (*s* = 0.1, Figures 2A and B) we see that increasing the rate of migration generally increases the probability of rescue in both the stepping stone as well as the island model. The reason is that for low costs of the rescue mutation we mainly see the positive effects of migration - demographic rescue and relaxed competition - whereas gene swamping can be neglected. The stepping stone and the island model show qualitatively similar patterns. The only difference is that for a given migration rate the positive effect of gene flow is larger in the island model as compared to the stepping stone model, and this difference increases with increasing number of demes. This is sensible because in the stepping stone model gene flow is restricted between neighboring demes, such that larger rates of gene flow are necessary to obtain the same positive effect of gene flow as compared to the island model.

**Figure 2:**
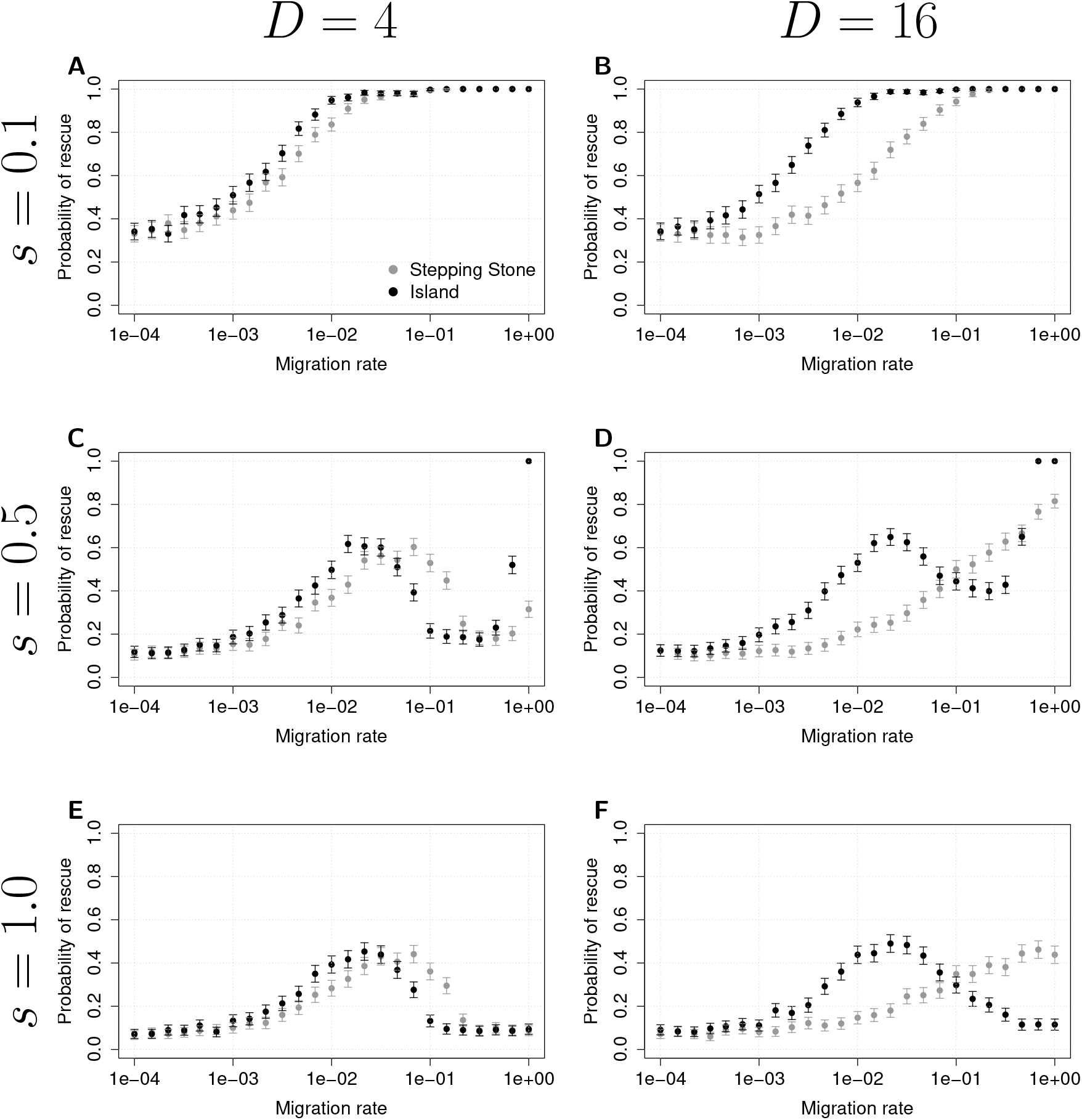
Comparison between stepping stone and island model. We show *P*_rescue_ as a function of migration rate for (A) *D* = 4, *s* = 0.1, (B) *D* = 16, *s* = 0.1 (C) *D* =4, *s* = 0.5, (D) *D* = 16, *s* = 0.5, (E) *D* = 4, *s* = 1.0, (F) *D* = 16, *s* =1.0. For all figures, *r* = 0.5, *z* = 0.02, and Θ = 4800.

Figures 2C and D show what happens when the cost of a mutation in the non-deteriorated environment is relatively high, but not lethal (*s* = 0.5). For *D* = 4 demes, we observe the effects of demographic rescue, gene swamping and relaxed competition. The interplay of demographic rescue and gene swamping causes a local maximum of the rescue probability for intermediate migration rates (around *m* = 0.01 − 0.1 in Figure 2C). Relaxed competition creates another increase in the probability of rescue when migration rates become for very large (around *m* > 0.5 in Figure 2C). As before, the island and the stepping stone model appear qualitatively similar for four demes. However, when the number of demes increases (*D* = 16 in Figure 2D) the two models show qualitatively different behavior. In the stepping stone model the ranges where relaxed competition and gene swamping dominate start to overlap, such that the chance of evolutionary rescue monotonically increases with the migration rate (Figure 2D), whereas the two distinct peaks in the probability of rescue remain visible in the island model.

Finally, when rescue mutations are lethal in the old environment (*s* = 1), relaxed competition does not occur because lethal mutations do not survive until density regulation in the original habitat. We thus find that intermediate migration rates maximize the chances of survival (Figures 2E and F), which is due to two contrasting effects of demographic rescue and gene swamping [Uecker et al., 2014, Tomasini and Peischl, 2020]. Again, the main difference between the island and the stepping stone model is that the optimal migration rate for evolutionary rescue is larger in the stepping stone model than in the island model, and this difference increases with increasing number of demes.

### Effective migration rate between habitats can explain differences between models

Gene swamping and relaxed competition are modulated by the amount of migration between different types of habitats, rather than the rate of migration between different demes. Roughly speaking, migration between demes that share the same environmental conditions does not affect the probability of establishment of a rescue mutation and has very little impact on local population sizes. We thus define an “effective migration rate” 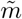 for the stepping stone model that quantifies the rate at which individuals migrate between the non-deteriorated and the deteriorated habitats. While this effective migration rate is difficult to calculate precisely, we present a simple heuristic derivation in Appendix A that shows that the effective migration rate between habitats in the stepping stone model is 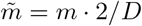.

We find that our heuristic effective migration rate can indeed explain the differences between the stepping stone model and the island model in most cases (Figure 3). Only when the cost of the rescue mutation is intermediate (*s* = 0.5) and the number of demes is large (*D* = 16) do we see notable qualitative differences between the two models. Apparently the reason for this is that while the effective migration rate captures the effects of demographic rescue and gene swamping quite well, relaxed competition depends on the migration rate in a more complex way in the stepping stone model. Indeed, when we perform simulations where we reduce the effect of relaxed competitions via Beverton-Holt density regulation [Beverton and Holt, 1957] with low growth rates, the discrepancy between the stepping stone model and the island model tends to disappear (see figure S5 in the supplemental material).

**Figure 3:**
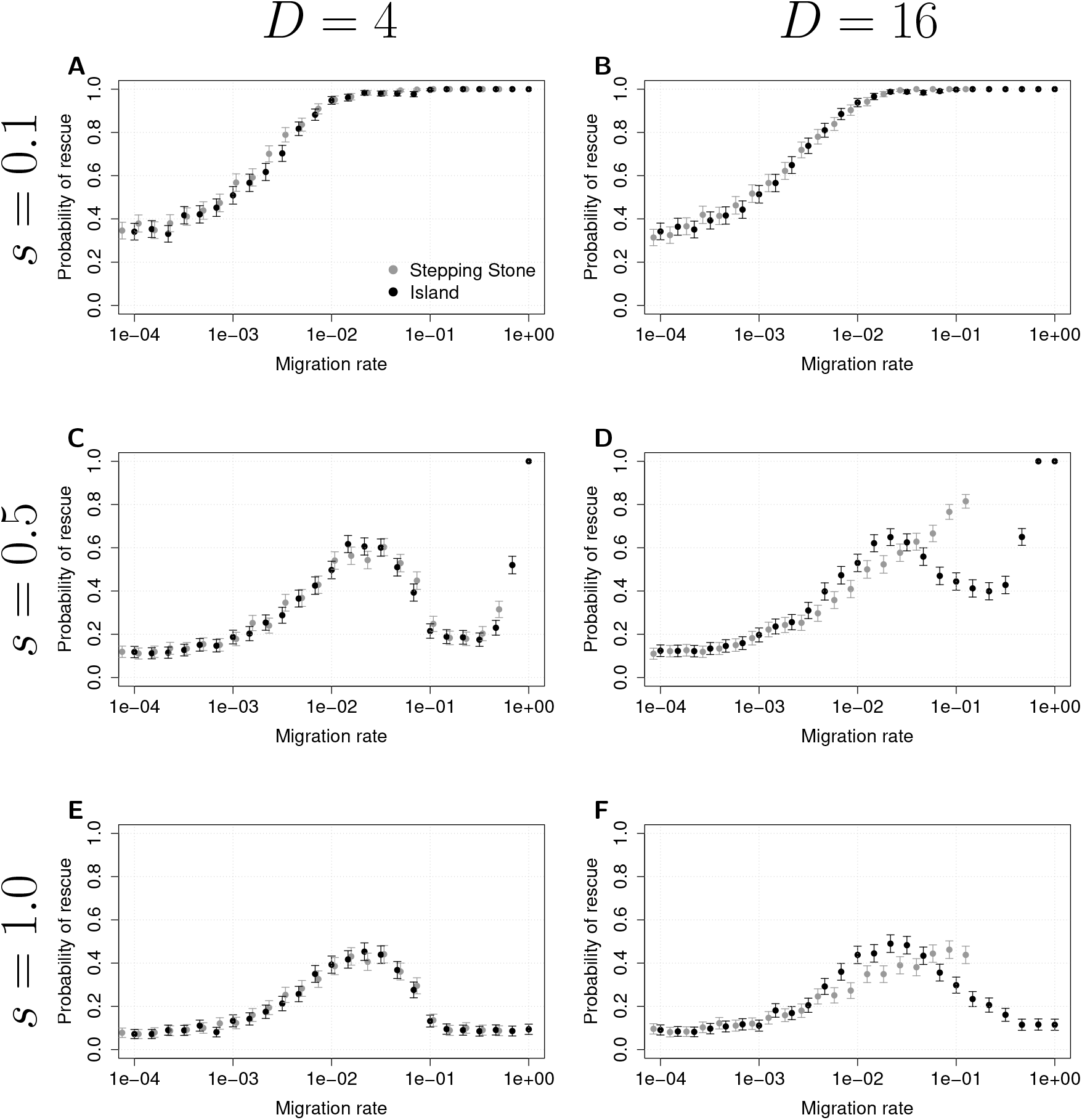
Probability of evolutionary rescue as a function of the “effective migration rate”. The data are the same as in figure 2, but we adjusted the migration rate in the stepping stone model, *m*_S_, with respect to the migration rate in the island model, *m*_I_, as *m*_I_ = *m*_S_ · 2/*D*. Chosen examples are the same as in figure 2: (A) *D* = 4, *s* = 0.1, (B) *D* = 16, *s* = 0.1 (C) *D* = 4, *s* = 0.5, (D) *D* = 16, *s* = 0.5, (E) *D* = 4, *s* = 1.0, (F) *D* = 16, *s* = 1.0. For all figures, *r* = 0.5, *z* = 0.02, and Θ = 4800.

While strong qualitative differences between the two models can be found only for intermediate values of *s*, we notice nonetheless that the use of the effective migration rate is not perfect in the case of lethal mutations (*s* =1, see figure 3F), as the maximal probability of evolutionary rescue is slightly lower in the stepping stone model than in the island model. So far we have discussed the negative effects of gene flow only as they pertain to the removal of beneficial mutations from the deteriorated environment; however, gene flow has a secondary negative effect due to demography. While the net demographic effect is positive (because the carrying capacity in the non-deteriorated demes is maintained by density-dependence), this is the result of a balance between immigrants and emigrants. While the number of emigrants from the deteriorated area is very small for low *m*, it turns out that this negative effect is more important for high migration (see appendix A for a quantification of this effect). This explains why, when applying the effective migration rate to the stepping stone model, we observe a lower maximal probability of rescue.

### Results are not strongly affected by the choice of parameters

Figure 4 shows contour plots of the probability of evolutionary rescue *P*_rescue_ as a function of the migration rate as well as other parameters of the model; we picked a set of parameters for which we expect qualitative differences between the models (highly fragmented habitat, *D* = 16, and when not otherwise specified, *s* = 0.5). Figures 4A, C and E show the probability of rescue in the island model; we recover findings previously shown for evolutionary rescue in a two-deme model and in an island model [Uecker et al., 2014, Tomasini and Peischl, 2020] and show that intermediate migration rates are optimal for a large range of parameters. Rescue is more likely for slower environmental change *θ* (figure 4A), for low stress *r* (figure 4C), and for a mutation weakly selected against in the old environment (low *s*, figure 4E). We can also see the strong effect of relaxed competition for low *s* and high migration rates *m* (figure 4E). Uecker et al. [2014] have studied the effects of different parameters for the island model in further detail, thus here we will focus on differences with the stepping stone model.

**Figure 4:**
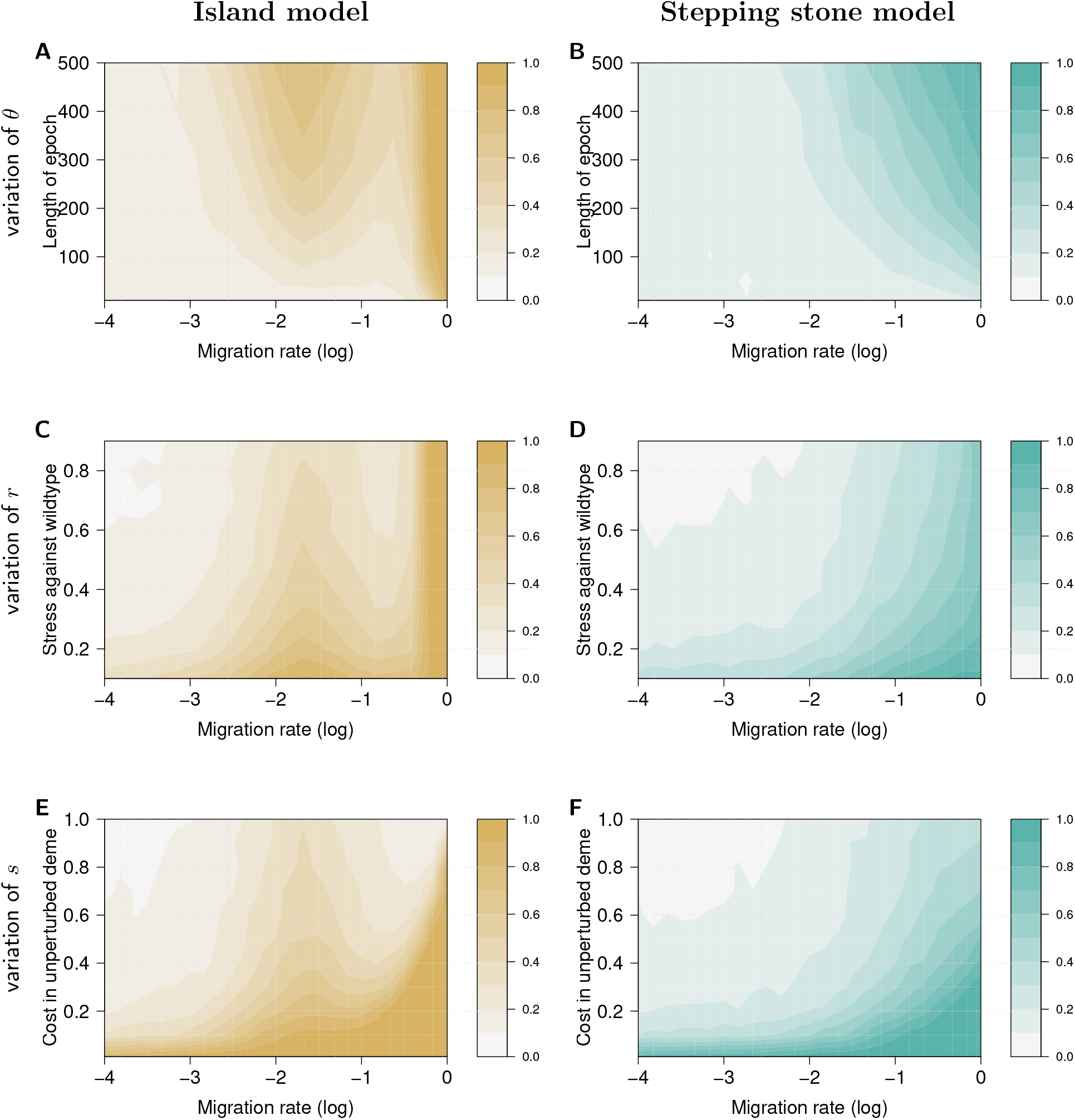
We show the contour plots of (A, C, E) the probability of rescue for the island model 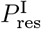, (B, D, F) the probability of rescue for the stepping stone model 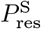, as a function of the migration rate *m*. We vary parameters one at a time: (A-B) the length of an epoch *θ*; (C-D) the harshness of environmental change r; (E-F) the cost *s* of the mutation in the unperturbed environment. Other parameters are *D* = 16, *z* = 0.02, and when not varying, *θ* = 200, *r* = 0.5 and *s* = 0.5.

Figures 4B, D and F show the contour plots for the probability of rescue in the stepping stone model. In contrast with the island model, these figures show that large migration rates tend to maximize the probability for rescue in the stepping stone model. In general, we observe patterns similar to those found in the island model, although for higher migration rates, as we would expect when considering the difference in effective migration rates between the models. Rescue is more likely for larger values of *θ*, and smaller values of *r* and *s*. We can see that the qualitative differences that we identified in the previous sections are robust for a wide range of parameters, in particular over the whole range of *r*, for 100 < *θ* < 500 and for 0.2 < *s* < 0.9. This suggests that the three cases that we studied in detail above (low *s*, intermediate *s* and high *s*) are representative of the majority of the range of parameters, with little change due to other parameters.

### Fragmentation can increase or decrease probability of rescue

Next we investigate the effect of increased habitat fragmentation in more detail. Since the effect of fragmentation is negligible in the island model [Uecker et al., 2014], we focus on the stepping stone model instead. By increasing fragmentation we mean increasing the number of demes over which a population is distributed in each run, while keeping the total population size (and the time it takes such that the whole habitat is deteriorated) constant. In other words, we split a large, well mixed population into an increasing number of smaller sub-populations. In figure 5 we look at the effect of fragmentation given different values of *m* and *s*. Figure 5 shows that in contrast to the island model, the number of demes can have quite dramatic effects on the probability of rescue. Only when gene flow is very low (yellow line in figures 5A, B and C, corresponding to *m* = 3.2 × 10^−4^) fragmentation has no effect on the probability for rescue. This is unsurprising, because with small migration populations evolve almost independently from one another. In such a scenario, evolutionary rescue depends on the total number of individuals in the habitat Ktot and not on how they are distributed over the environment. For larger values of gene flow (*m* =1 × 10^−2^ (orange line), *m* = 2.154 × 10^−2^ (magenta line), and *m* =1 × 10^−1^ (purple line) in Figure 5), we generally see that increasing fragmentation has negative effects on the rescue probability. The reason is simply that the positive demographic rescue effect of gene flow on evolutionary rescue becomes smaller with increasing number of demes (see also figures 2 and 3, and the derivation of the effective migration rate, Appendix A). When gene flow is very high (blue line, corresponding to *m* = 6.8129 × 10^−1^; for comparison, this is the second to last simulated point in figure 2), we generally see a rather strong increase in the rescue probability with increasing number of demes. This is due to a transition from gene swamping being dominant for high migration and low number of demes (where the effective migration rate between habitats is large) to relaxed competition being dominant if the number of demes is large (see also figures 2D and 3D). The only exception is when the cost of rescue mutation is low, where we find certain rescue in virtually all simulation runs if the migration rate is high (5A). The reason is simply that when a mutation is weakly selected against in the old environment, relaxed competition outweighs gene swamping across the whole range of values of the number of demes *D*.

**Figure 5:**
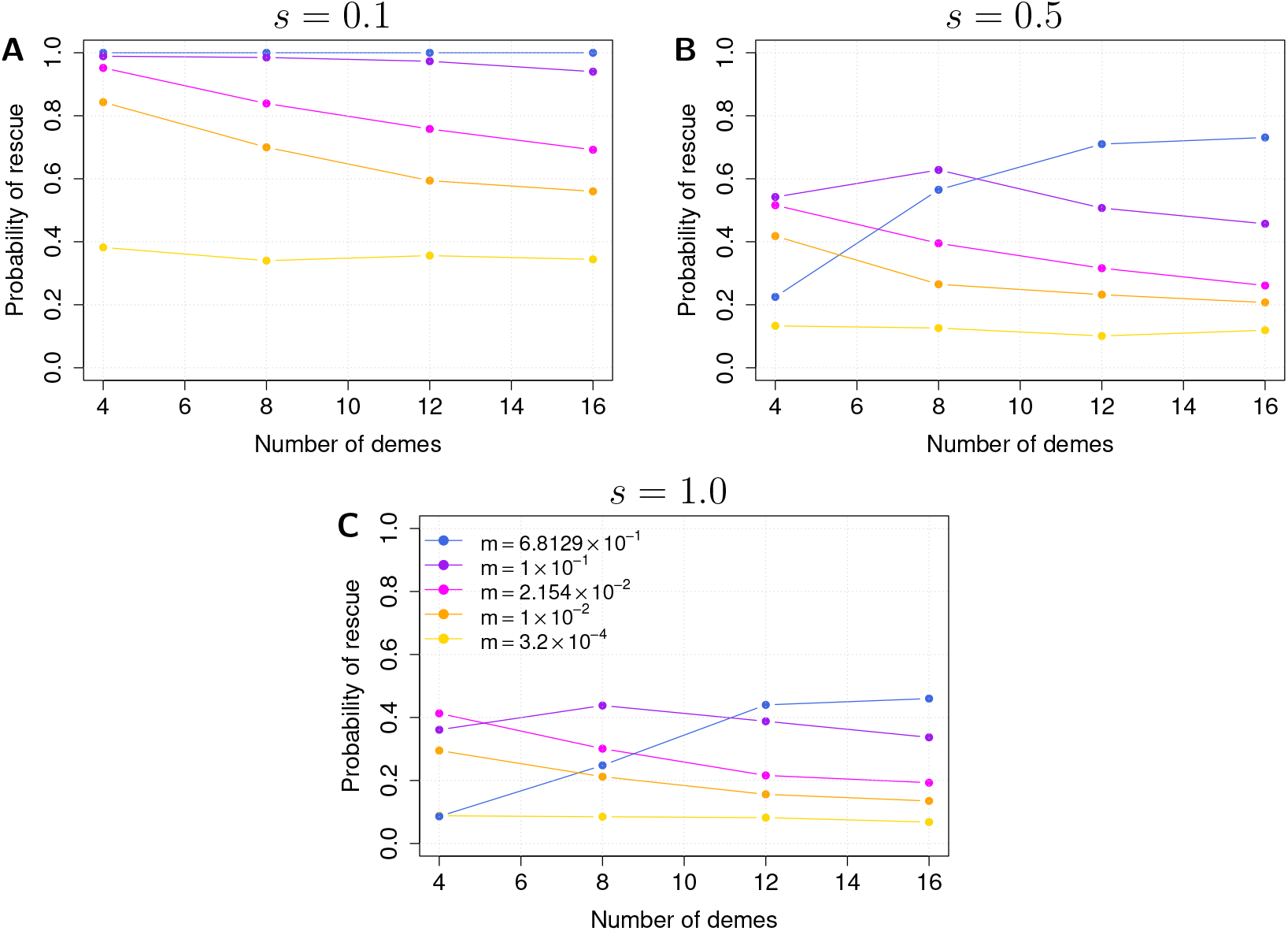
Probability of rescue as a function of the number of demes, for different values of migration rate and different values of *s*. For all figures, gold is *m* = 3.2 × 10^−4^, orange is *m* = 1 × 10^−2^, magenta is *m* = 2.154 × 10^−2^, purple is *m* = 1 × 10^−1^ and blue is *m* = 6.8129 × 10^−1^. (A) *s* = 0.1, (B) *s* = 0.5, (C) *s* = 1.0. In all figures, *r* = 0.5, *z* = 0.02, Θ = 4800.

### Random deterioration pattern

So far, we have considered evolutionary rescue in the island and the stepping stone models for scenarios where deterioration follows a spatial gradient (*i.e*. from deme 1 to deme *D*). While the order of deterioration of the demes is not relevant in the island model (because migration connects all the demes in the same way), it is important in the stepping stone model. This is once again because what determines the positive and negative effects of migration are the effective migration rates between types of environment, rather than between demes. In the stepping stone model with gradual deterioration, there is never more than *κm*/2 individuals moving between regions; to this effect, we can see the lastly deteriorated deme as a sort of valve that controls the flow of individuals between regions. If deterioration occurs randomly though, there will be more than just two demes exchanging migrants from one region to the other. In figure 6 we observe that when deterioration is random, island model and stepping stone model show very few differences. In particular, for mild mutation cost (*s* = 0.1, figures 6A and B) no difference at all can be noted. For cases with larger *s* we observe slight quantitative differences that are probably to attribute to slight differences in the flow of mutants between deteriorated and non-deteriorated area. In the island model, the local population in each deme depends on the environmental state of the deme (deteriorated or non-deteriorated, see *e.g*. figure S6E and F); however in the stepping stone model with random deterioration local density in each deme depends on whether the neighboring demes are in the same state, both in a different state, or split between non-deteriorated and deteriorated. The average effect on evolutionary rescue remains qualitatively the same.

**Figure 6:**
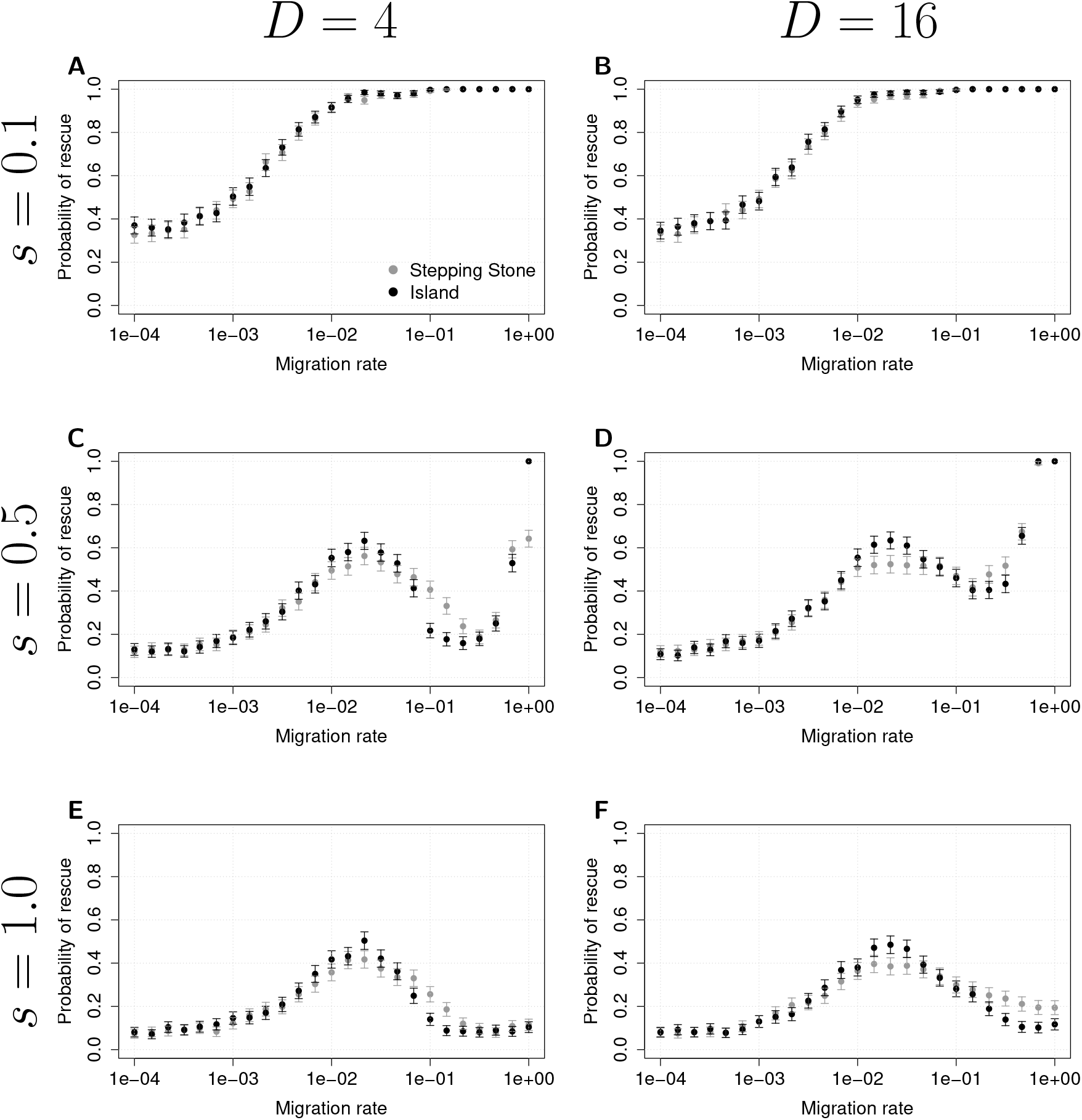
Comparison between stepping stone and island model with random deterioration. We show *P*_rescue_ as a function of migration rate for (A) *D* = 4, *s* = 0.1, (B) *D* = 16, *s* = 0.1 (C) *D* = 4, *s* = 0.5, (D) *D* = 16, *s* = 0.5, (E) *D* = 4, *s* = 1.0, (F) *D* = 16, *s* = 1.0. For all figures, *r* = 0.5, *z* = 0.02, and Θ = 4800.

## Discussion

In this paper we analyzed how the interplay between spatial structure, gene flow and fragmentation affects evolutionary rescue. We compared the island model and the stepping stone model with environmental conditions deteriorating over space and time.

In the island model the number of demes in the population does not influence rescue (as shown previously by Uecker et al. [2014]), while in the stepping stone model fragmentation strongly affects evolutionary rescue (figures 2, 4). In both models, rescue is the result of the interaction of three previously identified effects of gene flow on adaptation and thus rescue [Uecker et al., 2014]: gene swamping [Bulmer, 1972, Lenormand, 2002], demographic rescue [Brown and Kodric-Brown, 1977], and relaxed competition [Uecker et al., 2014]. Gene swamping and demographic rescue depend mostly on the flow of individuals between the non-deteriorated and the deteriorated areas of the environment, while relaxed competition is also strongly affected by the cost *s* of the rescuing mutation in the old environment.

We showed that for very high or very low cost of the rescuing mutation *s*, the stepping stone model can be thought of as a two-deme model with individuals moving between deteriorated and non-deteriorated regions, and we defined an effective migration rate that allows to compare island model to stepping stone model (figure 3A, B, E, F). This approximation breaks down for intermediate cost *s* and high fragmentation *D* (figure 3D).

Our main result was to show that, ultimately, the spatial structure of the habitat (*i.e*. how demes are connected) matters only in function of how it regulates the flow of individuals moving between the deteriorated and the non-deteriorated region; in the island model, fragmentation of a habitat does not affect its connectivity, while in the stepping stone model, fragmentation decreases the flow between areas due to decrease of local population. As such, all of these models can be understood as two-deme models – models where what is important is the exchange between different environments within the habitat. As a direct consequence of this, theoretical insights of two-deme models (*e.g*. Tomasini and Peischl [2020]) – generally presenting less analytical complexity – can also provide insights for models with more complex spatial structure.

We also analyzed how fragmentation affects evolutionary rescue, and showed that the effect of fragmentation is highly dependent on gene flow *m* and cost of mutation *s* (figure 5). As expected, for low migration rates fragmentation has no effect, as each deme evolves almost independently from the other demes. For intermediate migration, we find that rescue tends to decrease with increasing fragmentation, because demographic rescue becomes less important as deme carrying capacity decreases. For high migration, rescue increases with increasing fragmentation, as relaxed competition becomes more and more dominant and gene swamping loses importance. When the mutation is weakly selected in the old environment, relaxed competition dominates for high migration rates, for any rate of fragmentation.

Finally, we showed that the two models are almost indistinguishable if demes are deteriorated randomly over space: this is due to the fact that with random deterioration, migration between deteriorated and non-deteriorated areas is not limited to the lastly deteriorated deme, but occurs on the interface of all the regions of the environment where two different environments are neighbors. Then, this is yet another case where the stepping stone model can be approximated by a two-deme model.

Because of the importance of gene flow between types of environment (as opposed to gene flow between demes), we argue that experimental set-ups should not ignore the interaction between migration rates and spatial structure; in particular, it is important to understand what happens when gene flow is modified directly (by the experimenter) or indirectly (by the effects of fragmentation). For example, studies such as Bell and Gonzalez [2011], analyzing the different rates of rescue events in an experimental island model or a stepping stone model, only consider spatial structure as an explanatory variable, and not the amount of gene flow – thus conflating the two. This may also help explain why some experimental set-ups have observed contrasting results about the net effect of gene flow on evolutionary rescue [Carlson et al.,2014].

The fact that spatially explicit models can often be interpreted as two-demes model, if confirmed experimentally, could lead to improvement of conservation strategies of fragmented habitats. Our results suggest that knowledge about the exact spatial arrangement of sub-populations is not always required to understand rescue processes, instead one only needs to consider the rate at which gene flow occurs between different types of habitats.

When deterioration does not occur gradually in space (random deterioration), we show that spatial structure plays no role and evolutionary rescue occurs as in the island model (figure 6). This is because when demes deteriorate randomly, the exchange of individuals between the deteriorated and the non-deteriorated area occurs at many demes, as opposed to only one in the stepping stone model deteriorating gradually in space. This suggests also that evolutionary rescue in two or three spatial dimensions should look more like evolutionary rescue in an island model, even when migration is limited in space: this follows from the understanding that the deteriorated and non-deteriorated area are in contact over a larger “contact surface” than the one provided by the stepping stone model on a one-dimensional lattice. The exploration of different geometries in space, such as two- and three-dimensional lattices, could also yield a better understanding of the role of effects such as isolation by distance [Wright, 1943], as these geometries are arguably more general than one-dimensional lattices. However, the general conclusions of our study are not expected to change significantly in two- or three-dimensional habitats, as the effects we see (gene swamping, demographic rescue, relaxed competition) are not unique properties of a one-dimensional lattice.

We have assumed that migration is either between all demes, or only between neighboring demes. These are two extreme scenarios and an interesting extension would be to consider models where individuals disperse according to a distribution of migration distances. A better suited framework to explore such a theoretical model could be continuous space models (*e.g*. like in Barton [1987], Kirkpatrick and Peischl [2013]). For a narrow migration kernel, we expect to recover the behavior of the stepping stone model even in continuous space. As we increase the width of the kernel (*i.e*. the distance traveled by migrants) we would expect this model to be more similar to the island model. But since we do not expect gene flow to be isotropic in natural systems, because *e.g*. of physical barriers over the habitat and habitat range limits, it is difficult to predict whether chances of evolutionary rescue would increase or decrease when widening the migration kernel, due to the effects on evolutionary rescue when gene flow is not isotropic (see *e.g*. Kawecki and Holt [2002], Tomasini and Peischl [2020] for a discussion of asymmetric gene flow). In addition, Czuppon et al. [2021] studied the effects of biased migration on local adaptation and evolutionary rescue, using the same setting as in our current work. They show that dispersal schemes in which migration has a larger chance to occur towards deteriorated habitats increase the probability of rescue. This is in accordance to what we have found, as these migration schemes correspond to cases where the demographic positive effect of migration are stronger, and the negative effects are weaker.

In our model, density regulation determines how the probability of evolutionary rescue changes with high gene flow. When we choose a different mode of regulation (*e.g*. Beverton-Holt dynamics [Beverton and Holt, 1957]), the discrepancy between the island model and the stepping stone model disappears when accounting for the migration rate between environments (see figure S5 in the appendix). While we assumed that density is regulated only in the non-deteriorated environment, density regulation in the deteriorated environment is also expected to influence the chances of evolutionary rescue [Holt and Gomulkiewicz, 1997, Gomulkiewicz et al., 1999].

In our study, we considered single rescue mutations, thus ignoring multi-step rescue [Uecker and Hermisson, 2016] or the effect of the distribution of fitness effects and the fitness-landscape [Anciaux et al., 2018, Osmond et al., 2020]. It would be interesting to study the interactions of genetic architecture and spatial structure on evolutionary rescue, which would require including recombination rates and other effects (*e.g*. clonal interference, Gerrish and Lenski [1998]). Our results should however be a good approximation for populations that reproduce sexually with strong recombination, such that rescue mutations evolve independently. We do not expect our model to be predictive for organisms reproducing with low recombination rates if multiple rescue mutations are required for restoring positive growth rates, as we neglected competition between beneficial mutations [Gerrish and Lenski, 1998]. It has also been shown that the shape of the distribution of fitness effects as well as the recombination rate between loci harbouring rescue mutations will add complications that might interact with spatial structure [Uecker and Hermisson, 2016, Anciaux et al., 2018, Osmond et al., 2020]. In addition to genetic complications, we also neglected other important ecological factors such as competition (*e.g*. Osmond and de Mazancourt [2013], Klausmeier et al. [2020]).

Ultimately, our understanding of how dispersal interacts with evolution to save populations from extinction is still incomplete. While we analyzed the positive and negative effects due to how dispersal modifies the probability of rescue through demographic changes, we ignored several other eco-evolutionary feedbacks [Thompson and Fronhofer, 2019]. We also ignored the fact that dispersal rates can themselves be subjected to evolution, especially when a species’ range is shifting (*e.g*. Phillips and Shine [2006], Peischl and Gilbert [2020]). While such complexities are beyond the scope of the present study, they are paramount to reach a complete understanding of the feedback between evolution and dispersal, and how it affects the survival of species at risk of extinction.

Evolutionary rescue is still difficult to observe in natural populations [Carlson et al., 2014] with only a few recent putative examples (*e.g*. Epstein et al. [2016], Clayton and Spicer [2020], Muleta et al. [2021]). Direct comparison of our results to observed cases of evolutionary rescue in natural populations is therefore challenging. Laboratory model systems with automated dispersal show that gene flow can have different effects on evolutionary rescue, depending on whether dispersal is implemented locally or globally [Bell, 2013]. Such systems provide exciting opportunities to validate our theoretical results. Indirect evidence for our observation that antibiotic resistance should evolve more easily in populations with strong spatial structure comes from biofilm producing bacteria, in which antibiotic resistance is particularly widespread and difficult to control (see *e.g*. Vasudevan et al. [2019]). Furthermore, loss of flagellar function can confer a fitness advantage in the presence of an antibiotic in *Pseudomonas aeruginosa* [Rundell et al.,2020], suggesting further interactions between the evolution of antibiotic resistance and dispersal related traits. Our model could hence shed light on the difficulties to avoid the evolution of treatment resistance in spatially structured populations such as biofilms.

In conclusion, a better understanding of the role of spatial structure and the mode of dispersal would be helpful not only for conservation strategies of fragmented natural populations, but also in the context of public health and the evolution of antibiotic resistance.

## Authors contributions

MT and SP designed the project. MT and SP did analytical work. MT did all the simulations, figures, and wrote the first draft of the manuscript. MT and SP contributed to writing the manuscript.

## Acknowledgments

We thank Laurent Excoffier, Joachim Hermisson and Katie Peichel for stimulating discussions on this subject, and for helpful comments on the first manuscript. We also thank Lindi Wahl and Nick Barton for editorial work on an earlier version of this manuscript, as well as anonymous reviewers.

## Data archiving

The code used to perform simulations and generate figures, as well as the supporting information for this article can be found on github: https://github.com/mtomasini/EvoRescueFragmentation.

