## Supplemental material for "Evolutionary rescue in one dimensional stepping stone models"

### Evolutionary rescue in one dimensional stepping stone models – Supplemental Material

November 26, 2021

#### Appendix A: Demographic effects of migration

For all scopes and purposes, evolutionary rescue is affected by migration of individuals between demes in different states. This is because migration between two demes that are in the same state do not affect the probability of establishment of a beneficial mutation. In the main text we discuss the fact that there are two main effects due to gene flow, as outlined before in Uecker et al. [2014]: a negative effect due to the removal of beneficial mutations from the deteriorated environment, and a positive effect due to immigration of wild-type individuals in the deteriorated environment, leading to a temporary demographic rescue which slows down extinction. However, looking closely, demographic effects of gene flow can also be split into positive and negative. The positive effect is caused by the net immigration of individuals from the non-deteriorated to the deteriorated demes at every generation. The negative effect, due to the emigration of individuals from the deteriorated to the non-deteriorated part of the habitat. We now attempt to quantify these two effects.

#### Positive demographic effects of migration

Assuming that non-deteriorated are kept at carrying capacity by density regulation, in a two-deme model (equivalent to the island model) the total number of immigrants in the deteriorated area is

$$N_{\text{IM}}^{(\text{in})} = \kappa_{\text{IM}} \frac{m_{\text{IM}}}{2} \,. \tag{1}$$

where $\kappa_{\text{IM}}$ is the carrying capacity of the non-deteriorated deme. Here we use the subscripts IM to indicate that these are the quantities that come into play to determine rescue in an island model. While in an island model $\kappa_{\text{IM}}$ shrinks in time, on average its effect will be the same as having only two demes with $\kappa_{\text{IM}} = K_{\text{tot}}/2$, where $K_{\text{tot}}$ is the number of individuals over the whole habitat (see table 1 in the main text). To obtain the same net positive effect in the stepping stone model, we require

$$N_{\text{IM}}^{(\text{in})} = N_{\text{SSM}}^{(\text{in})} = \kappa \frac{m_{\text{SSM}}}{2} \tag{2}$$

If we remember that $\kappa = K_{\text{tot}}/D$, and we equate equation (1) to (2), we find that

$$\frac{K_{\text{tot}}}{D} \frac{m_{\text{SSM}}}{2} = \frac{K_{\text{tot}}}{2} \frac{m_{\text{IM}}}{2} \,, \tag{3}$$

and thus,

$$\boxed{m_{\text{SSM}} = m_{\text{IM}} \frac{D}{2}} \,. \tag{4}$$

While equation (4) provides a good approximation for low to intermediate $m$, it is only valid as long as $m_{\text{IM}}^2 D^2/4 \ll 1$ (see next subsection and main text).

We know that there is a certain value of the migration rate $m$ for which positive effects of gene flow are maximized, and negative effects are minimized. The negative effects change linearly with $m$. In the island model, fragmentation does not affect this critical value, but in the stepping stone model it does:

for higher $D$, a larger migration rate is needed to obtain the right balance between positive and negative effects. Equation (4) allows us then to define the "effective migration rate".

#### Negative demographic effects of migration

As mentioned, the positive demographic effect comes with a slight negative demographic effect represented by the number of emigrants from the deteriorated deme. However, the net effect of demography will always be positive, as $\kappa$ will always be larger than the population in the lastly deteriorated deme.

##### Two demes / island model

The amount of emigrants in two demes (equivalent to the loss in the island model) is easily calculated. After a few generations after a deme is deteriorated, the whole habitat reaches a state of stationary demography. For two demes, the population in the deteriorated deme, $N_d$ follows

$$N_d[t+1] = N_d[t](1-m)(1-r) + \frac{m}{2}\kappa \,, \tag{5}$$

where population in the non-deteriorated deme is $\kappa$. At equilibrium, roughly $N_d[t+1] = N_d[t]$, and the solution of (5) is

$$N_d = \frac{\kappa m}{2m + 2r - 2mr} \,. \tag{6}$$

##### Stepping stone model

We use a very rough approximation to calculate the number of individuals in each deme of the deteriorated region for the stepping stone model. Similarly to the case of two demes, after a few generations after a deterioration event, a demographic equilibrium is reached, where population in the lastly deteriorated deme is maintained at a certain level by incoming migration. This happens because the nearby original deme is filled at carrying capacity, and thus the input of migrants is constant. We have

$$N_i[t+1] = N_i[t](1-m)(1-r) + \frac{m}{2}(N_{i-1}[t] + N_{i+1}[t]) \,, \tag{7}$$

where $i$ denotes how distant a deme is from the first non-deteriorated deme, $i = 0$. For $i = 1$, we assume that $N_{i-1} = \kappa$. At equilibrium we just assume that $N_i[t+1] = N_i$. Then, we also assume that beyond a certain deme, the migrating population is so small to be negligible. For this example, we assume that $m/2mN_i \ll 1 \; \forall i \geq 4$, but the derivation is the same if we decided to set the limit further. However, neglecting all demes after $i = 4$ seems a sensible choice, given the population distribution shown *e.g.* in figure S6D). This is now a simple problem of substitution, solving first $N_3$, inserting it in the equation for $N_2$, and same until a form for $N_1$ is reached. Then,

$$N_1 = \frac{K\,m}{2\left(\frac{m^2}{4\left(\frac{m^2}{2\,(2\,m+2\,r-2\,m\,r)}+(m-1)\,(r-1)-1\right)} - (m-1)\,(r-1)+1\right)} \,. \tag{8}$$

#### Net difference of individuals

In the previous sections, we mapped the total demographic positive and negative effect due to gene flow. These two effects net a positive demographic effect, that will than interact with the negative effect due to loss of mutants to generate an ideal migration rate that maximizes the chance of evolutionary rescue. In figure S1 we show the positive demographic effect (calculated in the section *Positive demographic effects of migration*, solid line) as well as the net change in individuals in the island model (dotted line) and in the stepping stone model (dashed line). For the island model, the negative effects due to demography

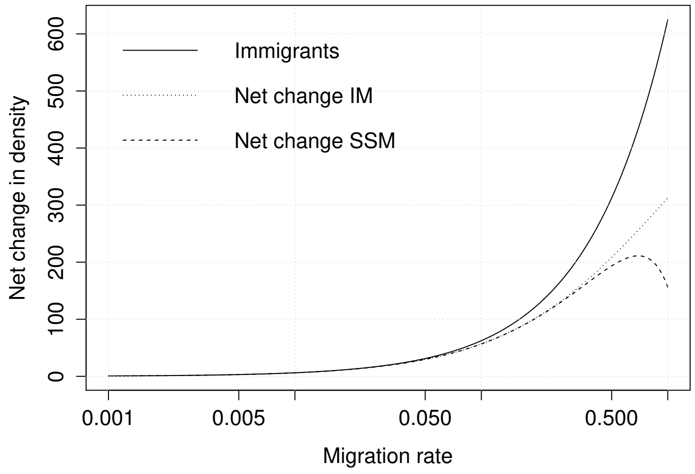


Figure S1: Net positive effect due to migration. Net change is $(\kappa - N_d)m/2$ for the island model, and $(\kappa - N_1)m/2$ for the stepping stone model (see text). Parameters: $D = 16$, $r = 0.5$.

are irrelevant in determining the optimal migration rate, as they are mostly negligible in that range.

But because with increasing $D$ the optimal migration rate increases in the stepping stone model, those negative effects will have an impact and decrease the maximal chance of evolutionary rescue with respect to the island model.

Thus, for the stepping stone model to obtain the same positive effect as the island model, the migration rate needs to be higher (by a coefficient $D/2$, see equation (4)). However, the negative demographic effects due to high migration impact the chances of evolutionary rescue, which will be slightly lower in that range. This explains the discrepancy in 3F when adjusting the effective migration rate.

Because these effects are strong only for very high $m$, we can still use formula (4) to map the stepping stone model into the island model for low to medium $D$ (when the optimal migration rate is not too high in the stepping stone model) and for low to intermediate $m$.

### Appendix B: Benchmark simulations

Details about how our simulations are performed can be found in the main text. We show that the simulation works for $m = 0$ (rescue occurring independently in $D$ demes, figure S2) and $\theta = 0$ (equivalent to one big deteriorated deme, figure S3). We also show the probability of rescue for our simulation with

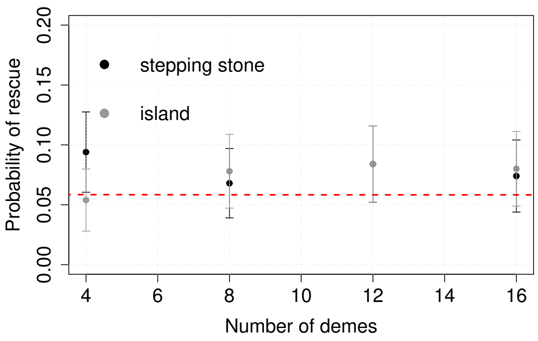


Figure S2: $P_{\text{rescue}}$ as a function of the number of demes in the island model (black) and the stepping stone model (gray), with $m = 1$. The red line is the expected value of evolutionary rescue for one large deme. Other parameters are: $s = 1$, $r = 0.5$, $z = 0.02$, $K_{\text{tot}} = 2 \times 10^4$, 500 replicates.

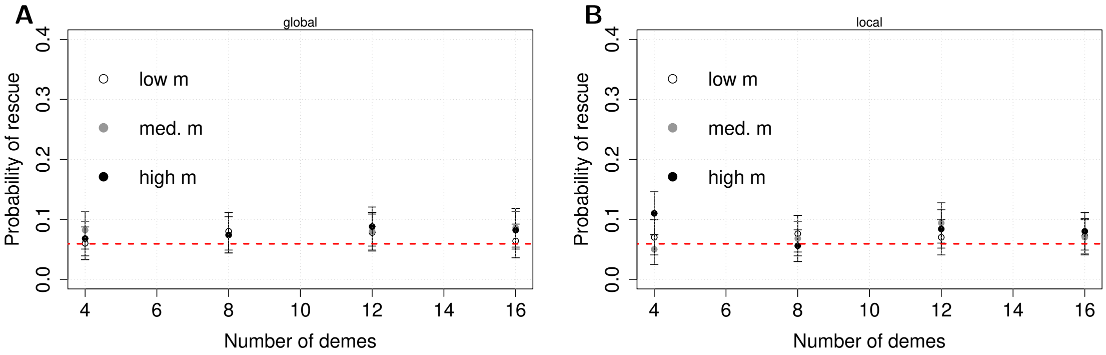


Figure S3: $P_{\text{rescue}}$ as a function of the number of demes in (A) the island model and (B) the stepping stone model, in the case of $\theta = 0$. The red line is the expected value of evolutionary rescue for $D$ independent demes. Low migration rate: $m = 3.2 \times 10^{-4}$; intermediate $m = 2.154 \times 10^{-2}$; high $m = 6.8129 \times 10^{-1}$. Other parameters are: $s = 1$, $r = 0.5$, $z = 0.02$, $\theta = 500$, 500 replicates.

only two demes (figure S4), compared to the approximation derived in Tomasini and Peischl [2020].

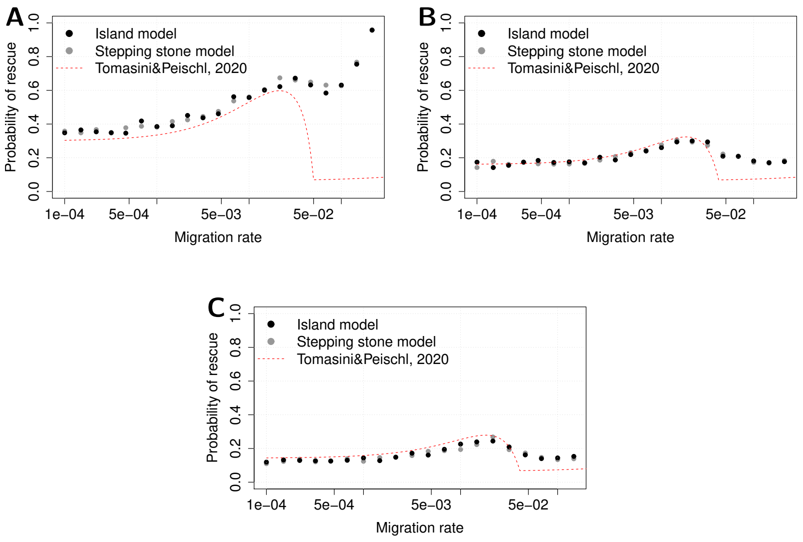


Figure S4: $P_{\text{rescue}}$ as a function of the migration rate with only two demes; (A) $s = 0.1$, (B) $s = 0.5$, (C) $s = 1.0$. The red line is the expected value of evolutionary rescue as calculated in Tomasini and Peischl [2020]. Other parameters are: $r = 0.3$, $z = 0.02$, $\theta = 500$, 1000 replicates.

### Appendix C: supplementary figures

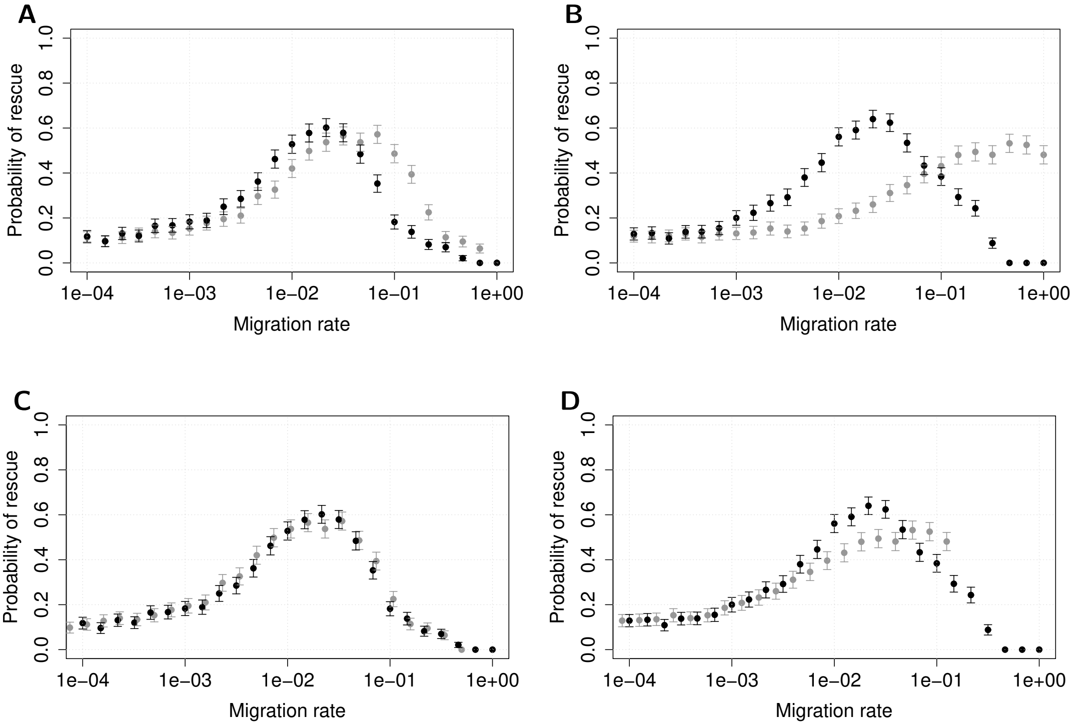


Figure S5: Comparison between stepping stone and island model. We show $P_{\text{rescue}}$ as a function of migration rate for (A, C) $D = 4$ and (B, D) $D = 16$, with density regulation according to Beverton-Holt dynamics Beverton and Holt [1957]. For all figures, $s = 0.5$, $r = 0.5$, $z = 0.02$, and $\Theta = 4800$, and Beverton-Holt growth rate $\rho = 1.5$.

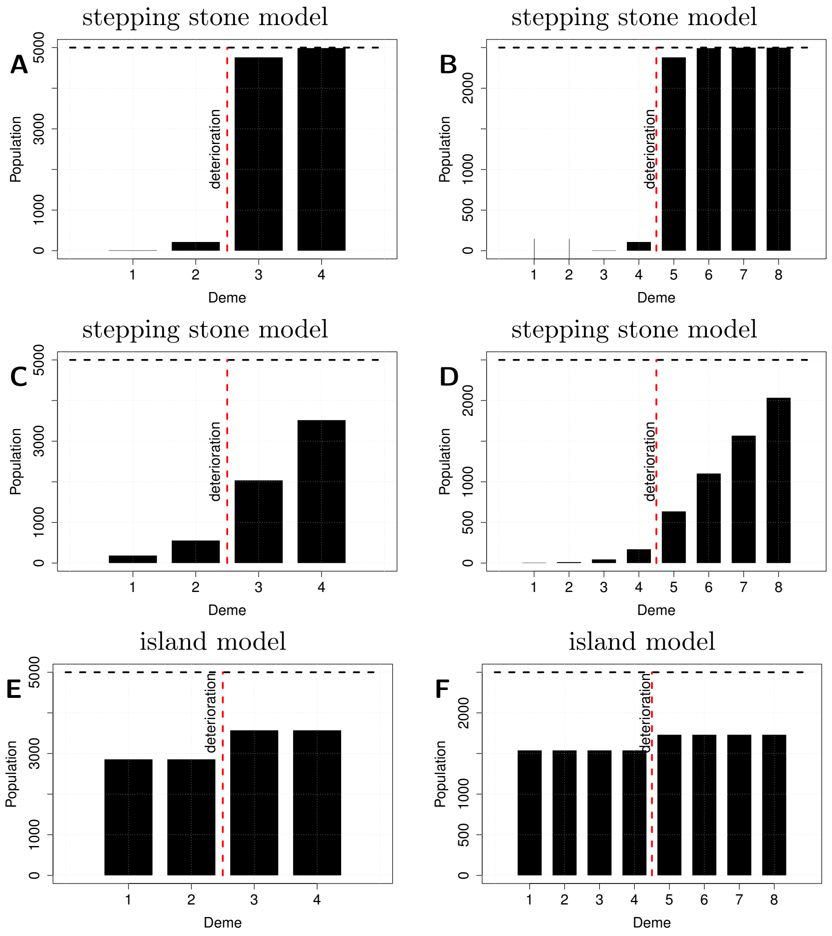


Figure S6: Deterministic populations densities before density regulation in demes at halfway of the habitat deterioration, for $D = 4$ (panels A, C and E) and for $D = 8$ (panels B, D and F). Panels A–D show the population in the stepping stone model, with panels A and B showing intermediate migration rate ($m = 0.1$) and panels C and D showing high migration rate ($m = 1$). Panels E and F show how population dynamics behave in the island model, with $m = 1$. Black dotted line is the carrying capacity per deme, red dotted line is the limit of the deteriorated area (left part). We can see how in the stepping stone model, demes are affected far away from the deterioration limit when migration is higher, thus prompting much stronger relaxed competition. In the island model, all demes are affected in the same way depending on their environment. Other parameters are $K_{\text{tot}} = 20000$, and $r = 0.5$, and $\Theta = 4800$. Halfway is then $t = 2399$, (the generation before deterioration of the next deme).
